# Multi-compartment tumor organoids

**DOI:** 10.1101/2020.11.03.367334

**Authors:** Meng-Horng Lee, Yohan Suryo Rahmanto, Gabriella Russo, Pei-Hsun Wu, Daniele Gilkes, Ashley Kiemen, Tsutomu Miyamoto, Yu Yu, Mehran Habibi, Ie-Ming Shih, Tian-Li Wang, Denis Wirtz

## Abstract

Organoid cultures are widely used because they preserve many features of cancer cells *in vivo*. Here, we developed high-throughput oil-in-water droplet microtechnology to generate highly uniform, small-volume, multi-compartment organoids. Each organoid culture features a microenvironmental architecture that mimics both the basement membrane and stromal barriers. This matrix architecture, which allows accessing both proliferative and invasive features of cancer cells in a single platform, has profound effect on observed drug responsiveness and tumor progression that correlate well with *in vivo* and clinical outcomes. The method was tested on multiple types of cancer cells including primary cells and immortalized cell lines, and we determined our platform is suitable even for cells of poor organoid-forming ability. These new organoids also allow for direct orthotopic mouse implantation of cancer cells with unprecedented success.

## Introduction

Tumor organoids are widely used to maintain and study cancer cells isolated from patients^1–8^. For example, organoids have been used to identify molecular pathways that drive tumor progression^9–13^, discover potential cancer biomarkers^14–19^ and predict patient response to customized pharmacological treatments^1,7,8,20–22^. However, before the implementation of organoids at scale for reliable application in high-throughput drug screening and predictive modeling *ex vivo*, greatly enhanced consistency, parsimony, mimicry, and cell-seeding control are imperative^23–27,56^.

Here, we introduce a novel oil-in-water droplet microtechnology to produce a highly consistent, small-volume organoids culturing platform. The volume of each organoid culture requires 1μl instead of typically >50μl for conventional organoid cultures. Hence, for the same amount of tumor tissue, many more individual organoids can be simultaneously created. Unlike standard organoid culture, this parsimonious platform avoids having to move organoids/cells out of the original 3D extracellular matrix (ECM) and transfer them to multi-well assay plates before drug testing/screening applications, which greatly reduces organoid-to-organoid variability^1,8,22^.

Moreover, current organoid cultures are limited to the use of only one ECM component at a time, typically either Matrigel or collagen I. This is a major issue as cells in tumor organoids mostly grow (and invade/migrate little) in Matrigel, and mostly invade (and grow little) in collagen I^28–32^ (see also below). In other words, proliferation and invasion, two key drivers of malignancy, cannot be fully included simultaneously in conventional organoid cultures. These new organoids feature a two-compartment architecture composed of juxtaposed recombinant basement membrane (rBM) and main stromal protein collagen I. These new organoids allow us to study highly invasive cancer cells, which are notoriously difficult to encapsulate^33,34^. We demonstrate the predictive power of these new organoids in recapitulating tumor progression *in vivo* and drug responsiveness in patients. We also successfully implant our new organoids containing cancer cells that had never been successfully implanted in mice.

## RESULTS

### Control of organoid volume, matrix architecture, and tumor cell growth

During the initial steps of transformation and growth as a precursor lesion, carcinoma tumor cells are first confined by a basement membrane rich in extracellular-matrix (ECM) components laminin, collagen IV, nidogen, and heparan proteoglycans^35^. Then, through disruption of this basement membrane and via a switch from a proliferative to an invasive phenotype, tumor cells spread into the surrounding stromal matrix, which is rich in collagen I^36,37^. To mimic this complex ECM architecture, we have developed a novel oil-in-water droplet microtechnology that reliably and rapidly generates, small organoid cultures made of two matrix compartments (Fig. 1a). Briefly, we drop a liquid solution containing the first extracellular-matrix material (typically rBM Matrigel) into mineral oil. The surface tension between water and oil and the resulting Plateau-Rayleigh instability molds the matrix into uniform droplets of spherical shape. To add another ECM, we dispense a liquid solution containing the second ECM material to encompass the first sphere after gelation of the first ECM material. Additional compartments composed of different ECM materials can be generated by simply repeating this process (Fig. 1b). Below, we first present a organoid model in which cancer cells are enclosed inside an inner compartment composed of rBM Matrigel and an outer compartment of stromal ECM type I collagen and highlight how this more physiological architecture affects fundamentally tumor progression and drug responsiveness compared to conventional Matrigel-only and collagen-only organoids.

**Figure 1.**
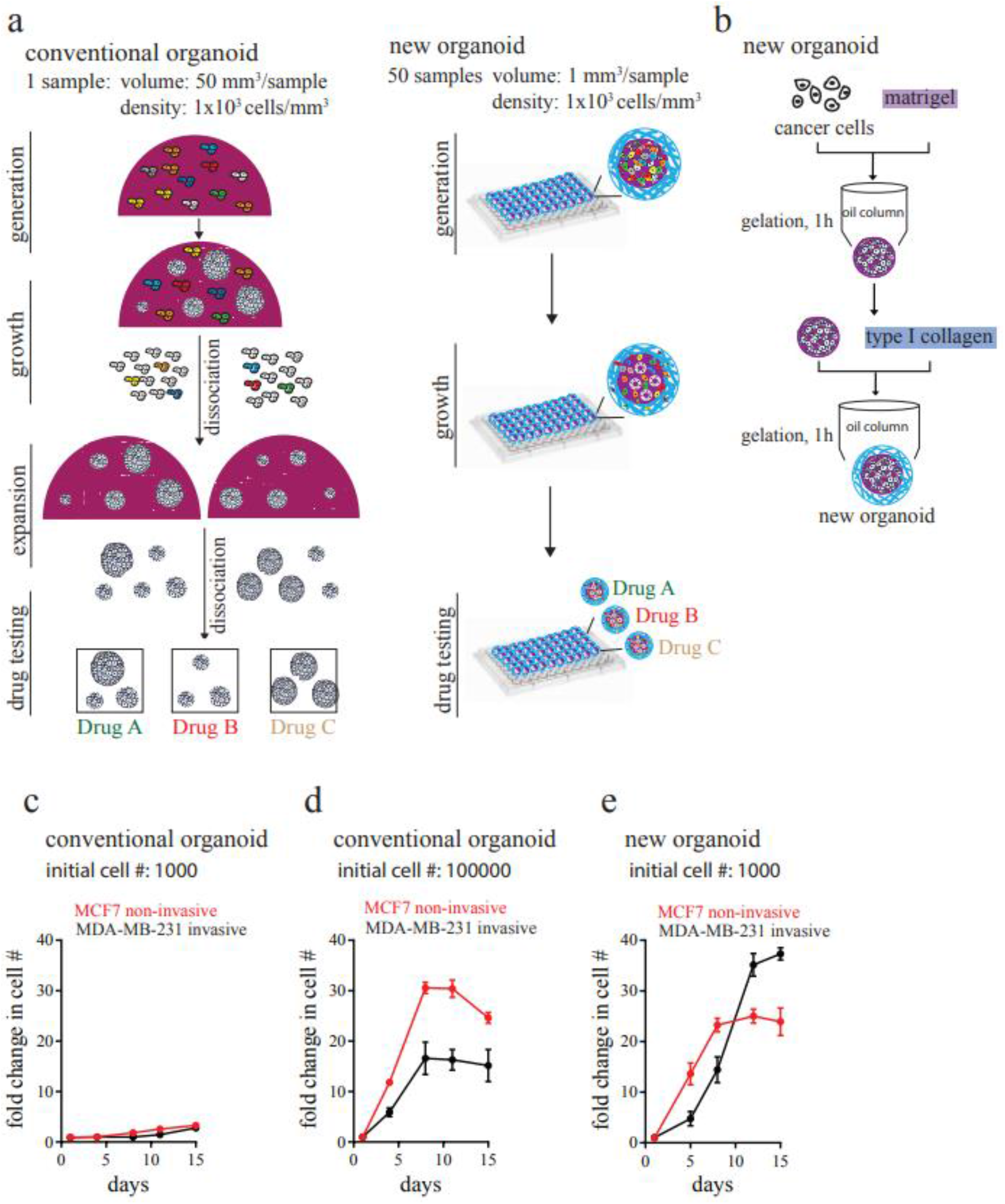
Oil-in-water droplet technology to make uniform, small-volume, two-compartment organoids. **a**, Schematics for the generation of conventional organoids *vs*. new two-compartment organoids and their use in drug-screening applications. In two-compartment organoids made via oil-in-water droplet microtechnology, the core contains cancer cells typically embedded in recombinant basement-membrane (rBM) matrix Matrigel; the outer compartment is made of type-I collagen. In conventional organoids, cancer cells are embedded in a single compartment, typically either Matrigel or type-I collagen. **b**, Design and manufacture of two-compartment organoids. **c**, and **d**, Growth rates of cancer cells in conventional organoids with different initial cell seeding numbers in Matrigel. **e,** Growth rates of cancer cells in two-compartment organoids. Cells used in panels c and d are MCF7 (red curves) and MDA-MB-231 breast cancer cells (black curves). In panels c-d, each curve encompasses three biological repeats and each repeat has five replicates for a total of 15 tested organoids.

Our organoid cultures are parsimonious (Fig. 1b). Our ability to generate 1μL Matrigel capsules allowed us to equally distribute cells into 50 cultures instead of just (typically) one when using conventional 50-μl organoid cultures. Thanks to this larger number of cultures, organoids do not have to be taken out of the original 3D matrix for cell expansion, as typically required for conventional organoid cultures.

Our oil-in-water microtechnology generates much more uniform organoid cultures. Controlling cell density is important because the density of cancer cells in 3D settings can influence cell phenotype and ability to migrate and invade^38,39^. The initial variation in cell numbers for the new organoids was <20% (Supplementary Fig. 1a). In contrast, if organoids were isolated from the 3D ECM and placed in multiple assay wells, the variation in cell numbers among wells was >80% because dissociated organoids have different sizes and contain different numbers of cells (Supplementary Fig. 1b-f).

Finally, the growth of cells can readily be optimized. Thanks to the small volume of the inner Matrigel compartment, we can create organoid cultures of highly controlled (initial) arbitrary cell density. By adjusting the seeding cell density, the growth of cancer cells in our organoid cultures can be readily optimized (Supplementary Fig. 2). In the new organoid culture, but not in conventional organoid cultures, cancer cells can successfully propagate even when the initial seeding cell number is very low (< 1000 cells) (Fig. 1 c and e). Moreover, invasive tumor cells in conventional organoid cultures expanded much more slowly than non-invasive cells (Fig. 1d). In contrast, invasive cells incorporated into the two-compartment organoids could propagate more than non-invasive cells (Fig. 1e). This is partly because the outer collagen compartment in the new organoid culture supported effective cancer cell invasion. As for conventional organoid culture, the progression of these new tumor organoids can be subjected to a battery of imaging assays (Supplementary Fig. 3).

### Two-compartment matrix architecture is necessary for cell growth and invasion

Our organoids can incorporate at least two different types of ECM compartments, which has important implications. We asked whether the order of these two ECM materials modulated the rate of proliferation and the mode of invasion of enclosed tumor cells. First, we generated four different types of organoids by choosing Matrigel and collagen I as ECMs for the inner and the outer compartments and studied the growth and invasive properties of breast cancer cells (Fig. 2a). As shown previously^40^, we found that cells in Matrigel formed larger tumors than in collagen I, confirming that the type of ECM used critically determined initial tumor size (Fig. 2b). Tumor cells in Matrigel expanded more rapidly than those cells in collagen I, independently of the composition of the outer compartment (Fig. 2c). Importantly, cells invaded the outer compartment as singlets and small aggregates only when it was collagen I (Fig. 2a). Tumors showed the fastest progression, measured by taking the ratio of the inner sphere volume to outer sphere volume, when the inner compartment was Matrigel and the outer compartment was collagen I (Fig. 2d). In contrast, the use of only one type of ECM material (i.e. the two compartments were made of either Matrigel or collagen I) significantly slowed down the progression of MDA-MB-231 tumors towards invasion and migration in collagen (Fig. 2e). The absence of either Matrigel or collagen I prevented tumor cells to grow and invade or migrate (Fig. 2f). All these phenomena were also observed for SUM149 and HCC1954 breast cancer cells (Supplementary Fig. 4). The reverse (non-physiological) architecture of an inner collagen core and an outer Matrigel compartment inhibited the progression of MDA-MB-231 cells (Fig. 2d).

**Figure 2.**
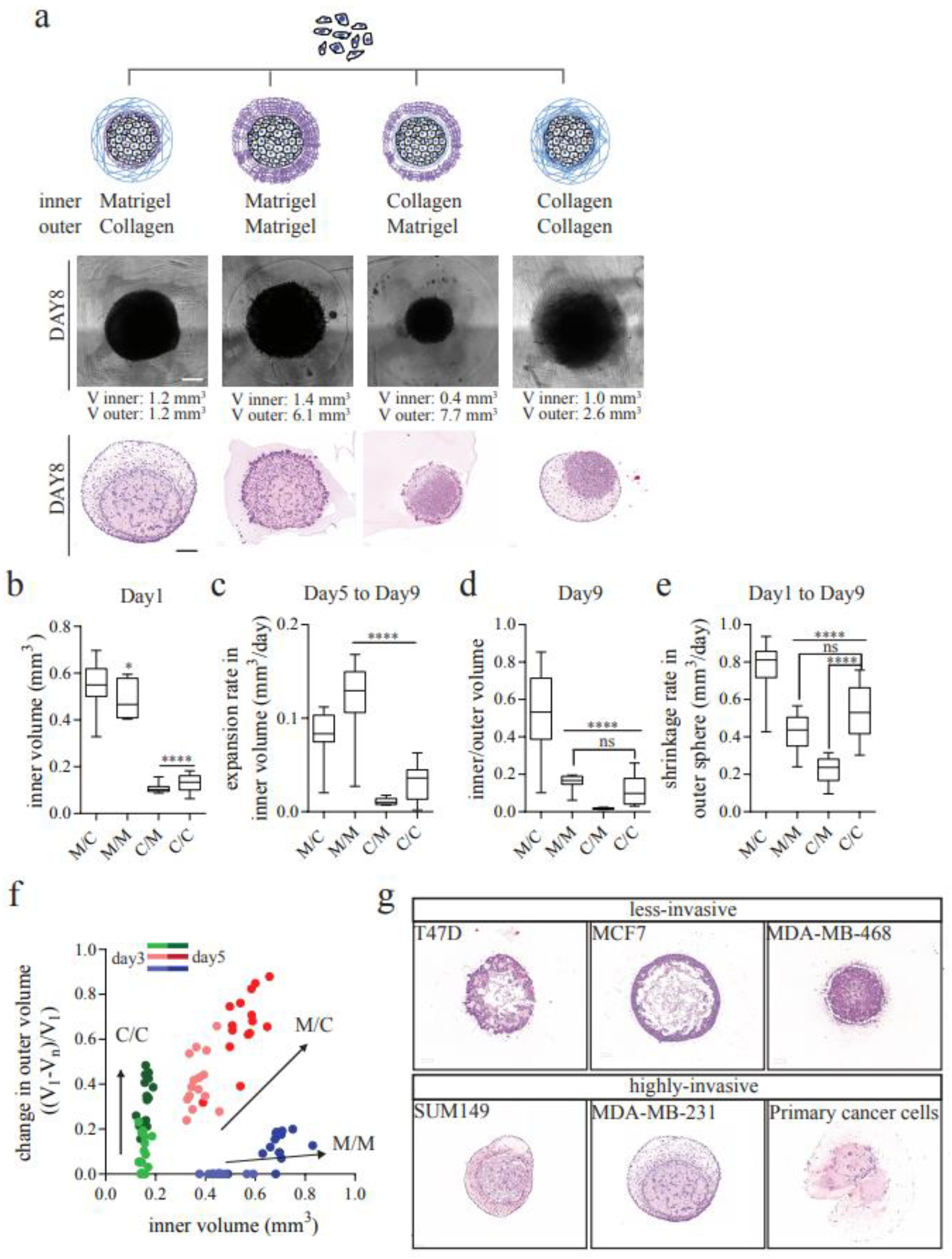
Differential progression of tumor cells using different configurations of matrix compartments. **a**, Representative phase-contrast images and H&E images of two-compartment organoids containing MDA-MB-231 breast cancer cells after 7 days in culture. **b,** Effect of ECM compartment composition for the growth of two-compartment organoids. Thanks to our oil-in-water droplet technology, the effect of the matrix composition of the inner core and outer compartment on drug responsiveness of cells could be examined. Bar graphs represent the mean ± SEM of the core volume of two-compartment organoids at day 1, **c**, their growth between days 5 and 9 in culture, **d**, their progression measured by the ratio of the volumes of the inner and outer compartments at day 9 in culture, and **e**, the shrinking rate of the outer compartment of new organoids in four different configurations of the two matrix compartments. The core and outer compartments either contain Matrigel (M) and/or collagen I (C) (**** p < 0.001). Note that C/C and M/M configurations are two-compartment organoids where inner and outer compartments are both made of collagen or both made of Matrigel, respectively, which are equivalent to conventional organoids composed of a single compartment made of either collagen or Matrigel only. In panels b-e, parameters for each organoids configuration encompasses three biological repeats and each repeat has five replicates for a total of 15 tested organoids. **f,** Shrinkage of the outer compartment and expansion of the core tumor compartment of new organoids between days 3 and 5. Each datapoint corresponds to the volumes of the inner and outer compartments of an organoid measured at day 3 (light color) and day 5 (dark color). **g,** Different patterns of tissue organization and invasion of new organoids composed of different types of cancer cells (cell lines and primary cells) and cultured for a week, as assessed by sectioning followed by H&E staining and imaging. Scale bar, 200 μm.

Together, these results indicate that the proper juxtaposed organization of the basement membrane-like compartment and collagen I compartment made possible by our new organoids, greatly impacts the progression and invasive properties of carcinoma cells. With a Matrigel/collagen bi-layered architecture, cancer cells in the new organoids can progress via proliferation, invasion and remodeling of the surrounding ECM. Depending on the type of cancer cells used and diverse cell-ECM and cell-cell interactions controlled by the modulable architecture of these new organoids, cells showed unique ways to organize themselves and the ECM (Fig. 2g).

### Predicting drug responsiveness using two-compartment organoids

Next, we show that drug responsiveness of cancer cells in our new culturing platform correlates with predicted clinical drug responsiveness. Clinical investigations have shown that tumors presenting high estrogen receptor alpha (ER) expression are associated with better response to hormone therapy and poorer response to chemotherapy^41–43^. The breast cancer cells grown in our culturing platform correctly predicted that a low dose of the estrogen receptor modulator tamoxifen, while slightly promoting the growth of ER^-^ MDA-MB-231 cells, could inhibit the growth of ER^+^ MCF7 cells (Fig. 3b and c). In contrast, the same cells grown using conventional organoid culture predicted erroneously the same response of these ER^+^ and ER^-^ cells to tamoxifen treatment (Fig. 3g and h). Next, the new organoids containing ER^+^ cells correctly showed no response to treatment with the cytotoxic drug paclitaxel (Fig. 3d and e). The new organoids also correctly predicted the response of ER^-^ cells to paclitaxel. Using the conventional method to grow cancer cells again erroneously predicted the same response of MCF7 and MDA-MB-231 cells to paclitaxel (Fig. 3i and j).

**Figure 3.**
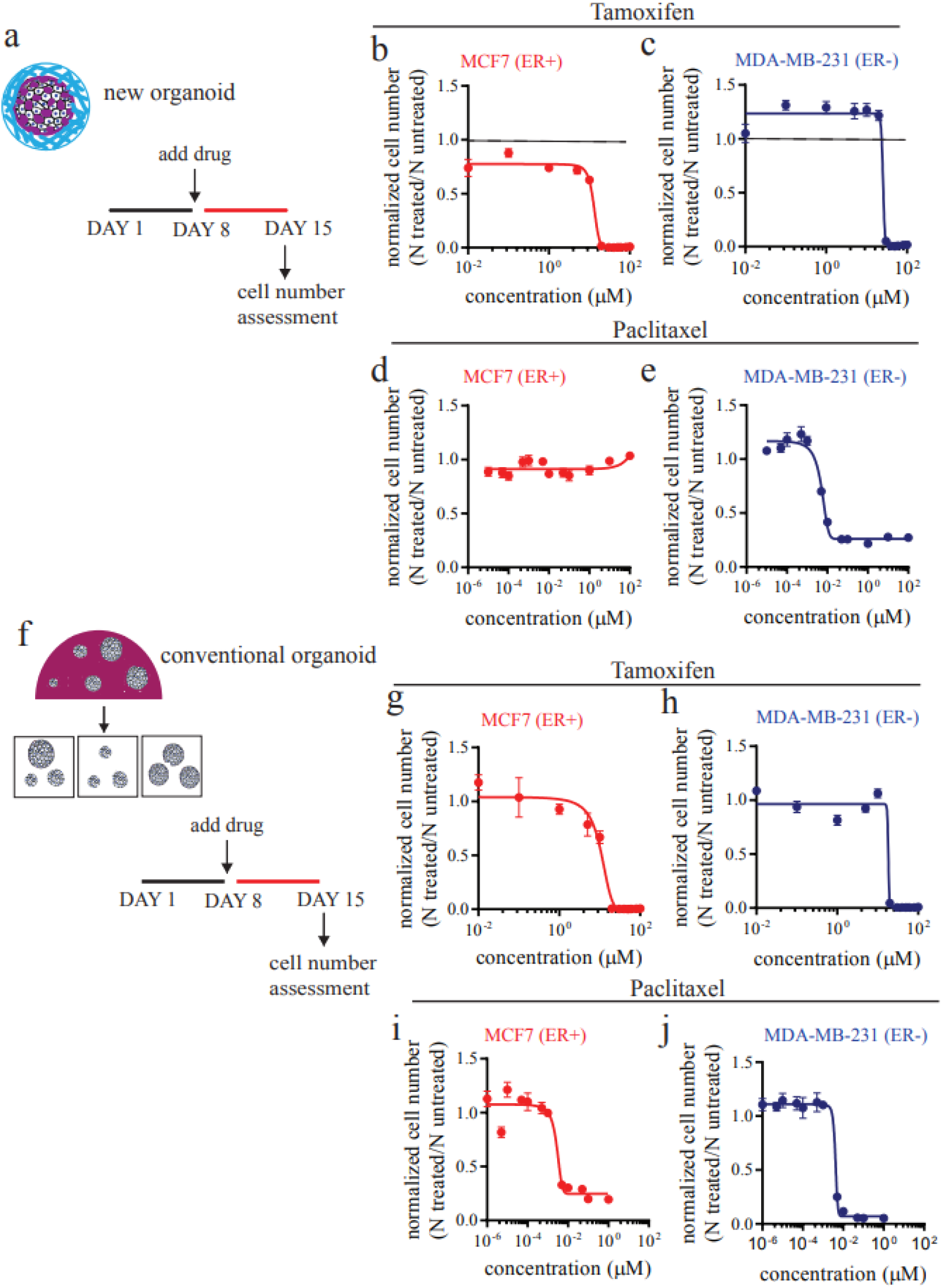
Drug responsiveness of conventional and two-compartment tumor organoids. **a-j**, Two-compartment organoids and conventional organoids containing estrogen receptor-positive (ER+) MCF7 cells or ER^-^ MDA-MB-231 breast cancer cells and treated with ER modulator tamoxifen or cytotoxic drug paclitaxel. Responsiveness of MCF7 breast cancer cells (**b, d, g** and **i**) and MDA-MB-231 breast cancer cells (**c, e, h** and **j**) incorporated into the Matrigel core of (**b-e**) two-compartment organoids and (**g-j**) conventional organoids to (**b, c, g** and **h**) tamoxifen and (**d, e, i** and **j**) paclitaxel. For the two-compartment organoids, the outer compartment is made of collagen I. Drug doses are indicated in the panels. Each measurement in panels b-e ad g-j encompasses three biological repeats and each repeat, and each repeat has five replicates for a total of 15 tested organoids.

### Validation of new organoids *in vivo*

Next, we compared the progression of the new organoids *in vitro* and *in vivo* in three different cancer models. First, we analyzed the progression of primary cells isolated from the doxycycline-inducible *Arid1a* and Pten conditioned knockout mice in our two-compartment matrix to the progression of the cells in orthotopically implanted tumors in mice (Fig. 4a). *ARID1A* and *PTEN* are two key tumor suppressors in endometrioid carcinoma, the most common type of human uterine carcinoma and ARID1A mutation and loss of its expression correlate with tumor invasion in human endometrial carcinoma ^44–48^. Deletion of *Arid1a* significantly accelerates tumor progression of *Pten*-deleted endometrial and ovarian carcinomas as evidenced by marked invasion and metastasis in genetically engineered mouse models ^49,50^. We then isolated the epithelial cells from the mouse uterus and cultured them in our organoid model. Untreated cells formed regular structures in the inner Matrigel compartment (Fig. 4b). After doxycycline treatment to delete *Arid1a* and *Pten*, cells started to propagate (Fig. 4c); the volume of the inner Matrigel sphere was reduced more rapidly (due to cell contractility) in treated cells compared to untreated ones (Fig. 4d). Accordingly, cell density was 20-fold higher in the doxycycline-treated organoids than untreated ones (Fig. 4e). Importantly, we observed cell invasion into the surrounding collagen matrix (Fig. 4f). A similar pattern of tumor progression was seen in mice which showed endometrioid carcinomas two weeks after *Arid1a*/*Pten* co-deletion by doxycycline ^49^. These carcinoma cells were highly invasive, with individual tumor cells infiltrating through uterine myometrium and permeating angiolymphatic spaces. In contrast, tumor formation was not observed in the absence of doxycycline administration (Supplementary Fig. 5).

**Figure 4.**
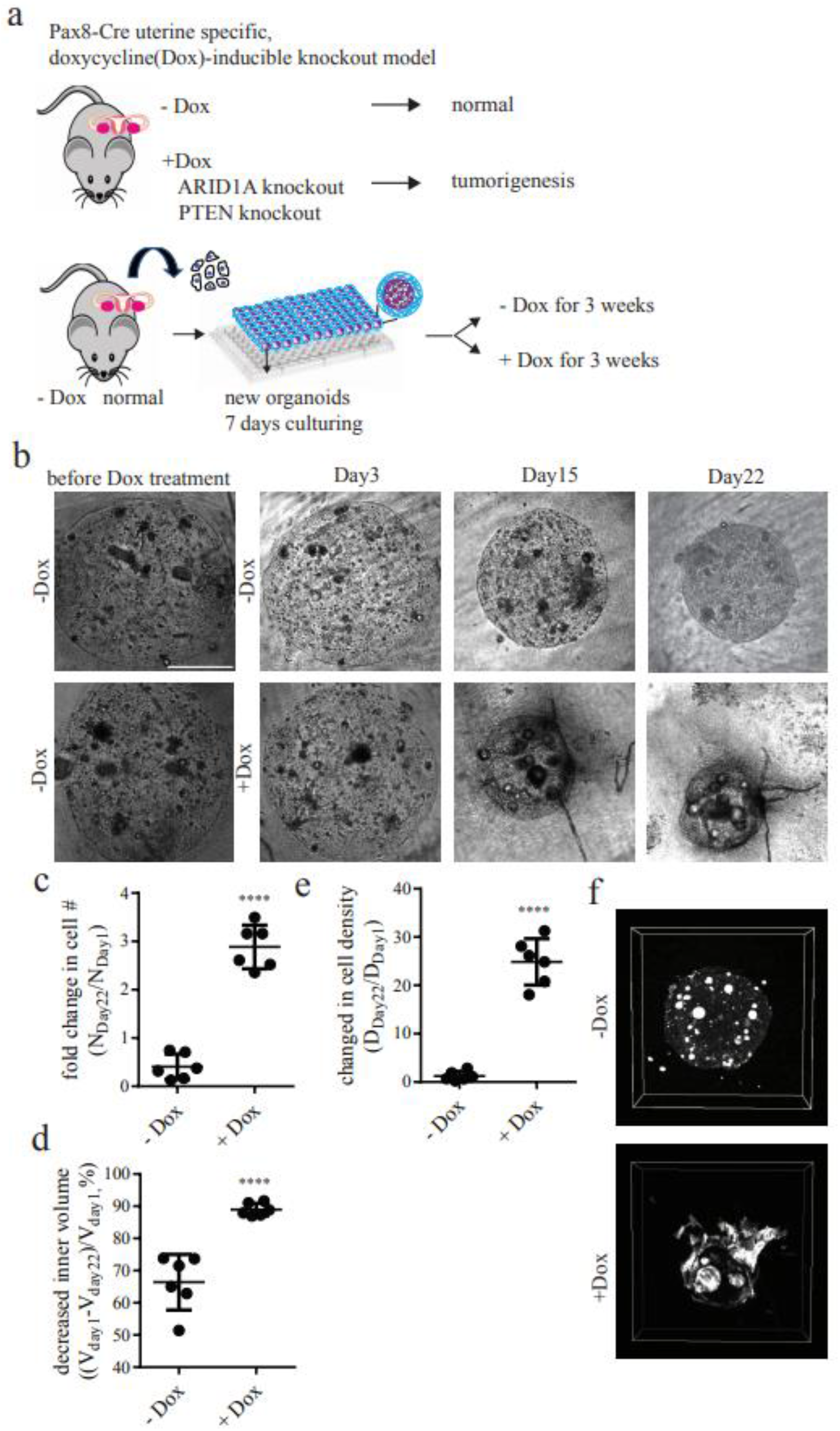
Two-compartment organoids mimic tumor progression *in vivo*. **a**, Schematic of two-compartment organoids containing primary cancer cells harvested from tumors that formed in doxycycline-inducible *Arid1a* and *Pten* knockout mice. **b**, Phase-contrast images of two-compartment organoids made of a Matrigel core containing the cancer cells and an outer collagen I compartment surrounding the core, in the absence and the presence of doxycycline. There is a significant increase in **c,** cell number **d**, volume shrinkage of the inner Matrigel sphere **e**, cell density after doxycycline treatment. **f**, Confocal images of new organoids in which the cells were labeled with F-actin. Scale bar, 200 μm. All two-compartment organoids had a Matrigel core and Collagen I as the outer compartment.

In a second model, we further assessed the predictive power of our organoids of outcomes *in vivo* by examining the effect of E-cadherin on tumor progression. High expression of E-cadherin is associated with the poor survival of patients diagnosed with basal breast cancer (Supplementary Fig. 6)^51^. To examine this scenario *in vitro*, we chose MDA-MB-231 cells, a commonly used basal type cell line that does not express E-cadherin. We employed a gain-of-function approach by supplying E-cadherin exogenously to MDA-MB-231 cells, termed here MDA+Ecad cells^52^. In our organoids, before reaching the same tumor size, the progression rate of MDA+Ecad tumors was twice as high as control MDA-MB-231 tumors (Fig. 5a). After two days, MDA+Ecad tumors entered a growth phase, while it took nine days for control MDA-MB-231 tumors to reach a growth phase (Fig. 5b and c). When cells were orthotopically implanted in the mammary fat pad of female NOD/SCID mice, their growth in the xenograft model correlated closely with our *in vitro* results using the new organoids: MDA+Ecad tumors displayed a two-fold faster growth than MDA-MB-231 tumors (Fig. 5d). Within five weeks, the MDA+Ecad tumors reached a size of 1 cm^3^, while it took 10 weeks for MDA-MB-231 tumors to reach the same size. MDA+Ecad tumors showed a significantly shorter lag phase comparing to the control MDA-MB-231 tumors (Fig. 5e and f), all results predicted by the new organoids.

**Figure 5.**
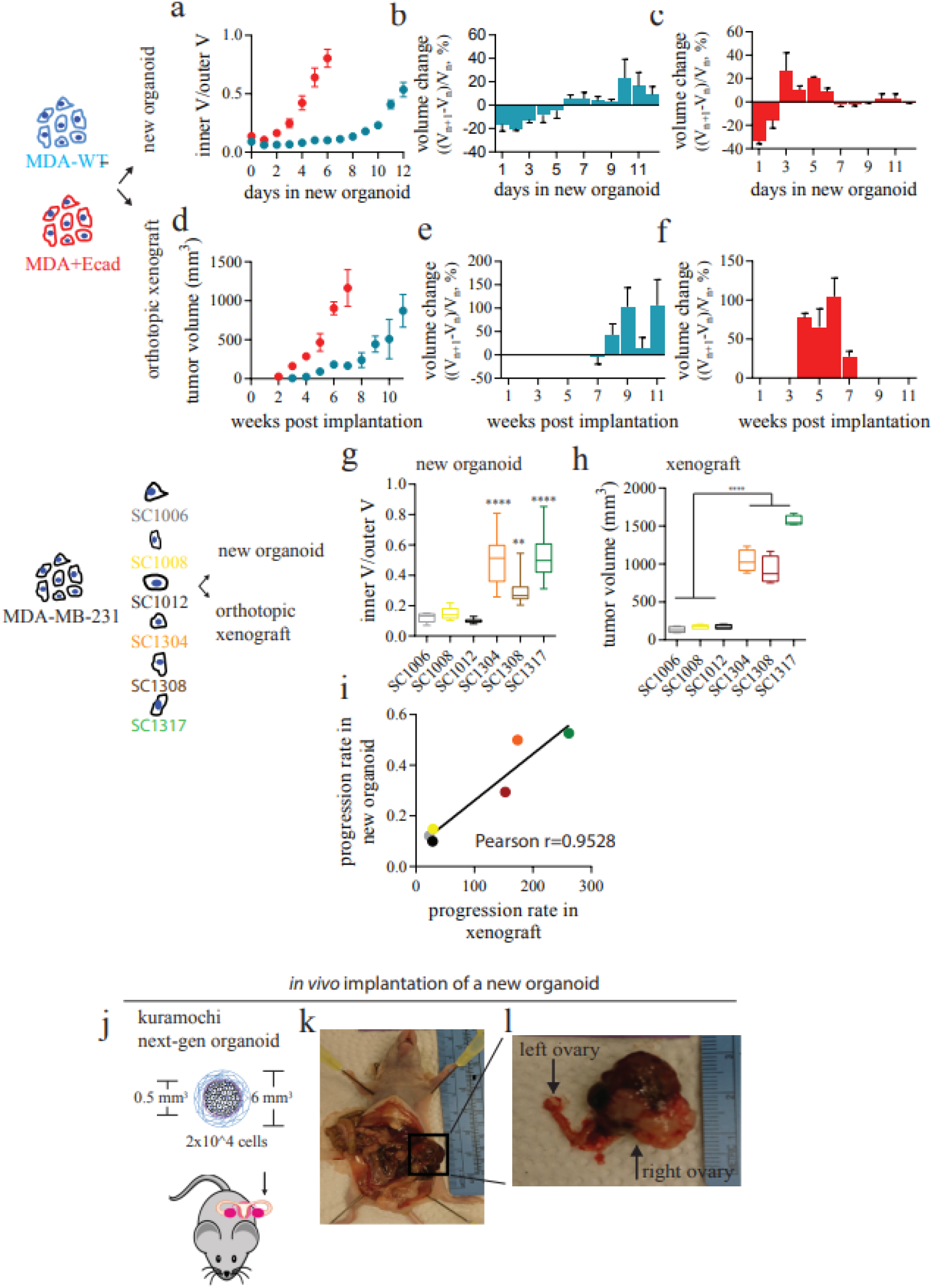
Tumor growth in two-compartment organoids correlates with tumor growth *in vivo*. Tumor growth in **a**, the two-compartment organoids *in vitro* or **d**, the mouse mammary fat pad *in vivo*. Growth patterns of (**b**, **e**) MDA-MB-231 scramble control breast cancer cells (MDA-WT) and (**c, f)** MDA-MB-231 cells expressing E-cadherin (MDA+Ecad) in (**b**, **c**) the two-compartment organoids and (**e**, **f**) in the mouse mammary fat pad of NOD/SCID mice. The bar graphs represent the mean ± SEM of the tumor volume of organoids containing different single-cell clones selected from parental MDA-MB-231 cells. These different single-cell clone cells were either **g**, incorporated in the two-compartment organoid or **h**, implanted into the mouse mammary fat pad. **i,** Correlation between tumor progression in the new organoids and in orthotopic xenograft mice. **j,** Orthotopic implantation of two-compartment organoids into the ovary in the female nude mice. These organoids contain high-grade serous ovarian cancer Kuramochi cells in their core. **k**, Tumor formation inside the abdominal cavity. **l**, Tumor formation in ovaries. All organoids used in Fig. 5 were composed of a Matrigel core containing the cancer cells and an outer collagen I compartment surrounding the core. The following numbers of measurements were conducted: five biological repeats and six replicates per experiment (panels a-c); two biological repeats and five replicates per repeat (panels d and e); three biological repeats and five replicates per repeat (panel f), and two biological repeats and four replicates per repeat (panel h).

To further confirm the correlation between outcomes predicted by modeling using the new organoids and *in vivo* outcomes, we next selected six single-cell clones (SCs) generated from the parental MDA-MB-231 cell line and subjected them to both modeling using new organoids and modeling using orthotopic xenografts *in* vivo. We correctly found that a subset of these clones displayed an aggressive growth, while a subset showed a low growth rate, both in the new organoids and in mice (Fig. 5 g-i; Pearson coefficient of tumor growth rates in two-compartment organoids and mice was 0.95).

Our new organoid culture can be used to generate orthotropic xenograft models. We implanted our two-compartment organoids containing high-grade serous ovarian cancer Kuramochi cells into the ovary of female nude mice (Fig. 5j). We chose this cell line because it is known to be non-tumorigenic and challenging - if not impossible - to grow in nude mice^53^. Using our two-compartment matrix as vehicle, Kuramochi cells grew and formed tumors in the implanted ovary (Fig. 5i). We did not observe tumors that randomly grew on other organs in the abdomen (Fig. 5k), indicating that using our new organoid to deliver the cancer cells can greatly facilitate their implantation and growth only on target organs.

## CONCLUSION

We have developed a novel method to produce 3D bi-layered tumor organoids that allows for high levels of consistency, control, and correlation to *in vivo* models of cancer cells, all while being high throughput. In addition to improving from traditional organoid culture techniques, our technique is also a translational system that can be used to study cancers *in vivo* that do not have well-established or feasible models for *in vivo* study.

We show that by utilizing oil-in-water droplet microtechnology, we can conveniently generate two-compartment organoids that better mimic the tumor microenvironment than more traditional single-ECM organoids. We have also demonstrated through various control experiments how vital the presence of both recombinant basement membrane and collagen layering is to generate a 3D *in vitro* model that closely correlates with *in vivo* data. The double-layered system allows for phenotyping and studying of cancer cell proliferation, invasion, and migration all in one experimental system and can be used to study cancers that are notoriously difficult to culture in the traditional 3D organoid systems, such as ovarian cancer. The novel technique we have developed allows for unmatched control of vital parameters to understanding how cancer cells proliferate and interact with the tumor microenvironment. The advantages of this system are extended to *in vivo* study of aggressive cancer types that lack well-established *in vivo* models. This approach will allow for further study of these cancers in pre-clinical mouse models to further improve patient outcome in the clinical setting. This platform will also allow for high-throughput drug screening, phenotyping, and predictive data acquisition that will greatly improve the too common disconnect observed between pre-clinical cancer research and clinical patient outcomes.

## Methods

### Two-compartment organoids made using oil-in-water droplet microtechnology

Unless specified, each two-compartment organoid was made by mixing a preset number of cancer cells with 1 μl Matrigel (Corning, Bedford, MA, USA). The mixture was dropped into mineral oil (Sigma-Aldrich) and incubated at 37°C for 30 min to allow for the gelation of Matrigel. The Matrigel sphere containing the cancer cells was harvested from the mineral oil and resuspended in a type I collagen solution (10 μl, Corning). This mixture was then dropped into mineral oil at 37 °C for 1 hour to allow for the gelation of type I collagen. The double-layered sphere was then collected. Each new organoid was cultured in suspension in a round-bottom well. In Fig. 4, we modulate the matrices used in the core and outer compartments to demonstrate the effect of organoid architecture on the drug responsiveness of cancer cells.

### Sample collection and tissue dissociation

Patients’ tumor samples were acquired under Johns Hopkins Medicine Institutional Review Board approval (IRB00164685). Fresh tumor tissues were kept in the tissue storage medium (Miltenyi Biotech) and then processed using the tumor dissociation kit (Miltenyi Biotech) and gentleMACS Octo Dissociator with Heaters (Miltenyi Biotech) according to the manufacturer’s instructions. Dissociated cells were washed and directly seeded into the new organoids.

### Time-lapsed and immunofluorescence microscopy

Time-lapsed images were collected every day for 1 week using a Nikon TE2000 microscope (Nikon) equipped with a 4x objective and a Cascade 1 K CCD camera (Roper Scientific). The images were automatically stitched using custom MATLAB software (The MathWorks).

For immunostaining, new organoids were fixed with 4% paraformaldehyde for 1 day and then incubated with the phalloidin (1:40; Invitrogen) and Hoechst 33342 (1:100; Invitrogen) for 1 day at 4 °C. For tissue clearing, the commercial kit (Visikol) was used according to the manufacturer’s instructions. Images of the stained cells were acquired with a Nikon A1 confocal microscope (Nikon) equipped with a 10× objective (Supplementary Fig. 7). The 3D images were reconstituted using NIS-Elements (Nikon).

### Drug response and cell viability

To examine the drug response of cancer cells, two-compartment organoids were manufactured, harvested and distributed in the wells of a 96-round-bottom plate. Each well contained a single organoid in 100 μl medium. Two times concentrated drug solution including tamoxifen (Sigma-Aldrich), paclitaxel (Selleckchem) as well as DMSO control 100 μl was added to each well. Cell viability was assayed using the cell viability reagent PrestoBlue (Thermo Fisher Scientific). The samples were incubated with PrestoBlue for 3 h. The fluorescence intensity of PrestoBlue was accessed using a Spectra Max plate reader (Molecular Device), according to the manufacturer’s instructions. Standard curves relating the controlled initial number of seeded cells in the organoids to the measured PrestoBlue fluorescence intensity was generated before data analysis (Supplementary Fig. 8).

### Histology and immunohistochemistry

All organoids were fixed with 10% formalin for 24 h and then processed by The Johns Hopkins University Oncology Tissue Services using a standard paraffin tissue embedding protocol and 4-μm sections were cut. For immunohistochemistry, formalin fixed paraffin-embedded sections were deparaffinized and rehydrated. Antigen retrieval was carried out using DAKO Target Retrieval Solution, equivalent to citrate buffer pH 6.0, or Trilogy, equivalent to neutral pH EDTA. Endogenous peroxidase activity was quenched in 3% H_2_O_2_. Sections were incubated with antibodies overnight at 4°C. Immunostains were visualized using DAKO EnVision+ System-HRP goat Anti-Rabbit IgG and DAKO DAB+ Substrate Chromogen System. Nuclei were visualized using hematoxylin counterstaining. Cover slides were mounted with Cytoseal 60.

To merge the images of two slides from the adjacent sections, we developed a customized MATLAB software. The nucleus-isolated images were rigidly registered in order to computationally align the cells in adjacent sections. Once aligned, the same registration method was applied to the antibody stain-isolated channels of the IHC images

### Animals

All animal studies were in accordance with animal protocols approved by the Johns Hopkins Medical Institute Animal Care and Use Committee. Generation of Arid1aflox/flox mice on the C57BL/6 background was described previously^49^. Briefly, Ptenflox/flox mice on the BALB/c background (Strain C;129S4-Ptentm1Hwu/J) were purchased from the Jackson Laboratory (Groszer et al., 2001). To express Cre recombinase specifically in the mouse uterine epithelium, we used Pax8-Cre mice which were generated by crossing mice expressing the reverse tetracycline-controlled transactivator (rtTA) under the control of the Pax8 promoter (Pax8-rtTA) with mice expressing Cre recombinase in a tetracycline-dependent manner (TetO-Cre). A knockout was initiated by treating mice with doxycycline either through oral gavage (2 mg/mouse/day) or subcutaneous implantation of doxycycline pellets (200 mg) when they reached puberty (6-8 week old).

### Orthotopic mammary fat pad injection

A detail procedure is provided elsewhere^54^. Briefly, 5- to 7-week-old female NOD/SCID mice were used. Mice were anesthetized and 1 × 10^6^ cells mixed with 100ml Matrigel (Corning) were injected into the mammary fat pad (MFP). Tumors were measured in three dimensions (*a, b*, and *c*), and their volume (*V*) was calculated as *V* = *abc* × 0.52.

### Orthotopic implantation of two-compartment organoids

Five- to 7-week-old female NOD/SCID mice were used. Once a mouse was anesthetized, we made an incision (7 mm) at the left back of the mouse to open the abdomen. The left ovary of the mouse was pulled out from the abdomen. We cut the surface of ovary and implanted a new organoid. Twenty μl of Matrigel was applied to secure the placement of new organoid on the ovary. After the implantation, we released the left ovary and let it back to the abdominal cavity. We closed the peritoneum by suture and closed skin incision using a stapler.

## Acknowledgements

We thank all members of the Wirtz Lab for discussions and feedback on this project. This work was supported through grants from the National Cancer Institute (U54CA143868) and the National Institute on Aging (U01AG060903) to D.W. and P.H.W.

## Contributions

M.H.L. and D.W. developed the hypothesis and designed experiments. M.H.L., G.C.R., D.G., performed experiments and data analysis. Y.S.R., T.M., Y.Y., T.L.W., and I.M.S. performed *in vivo* validation experiments and implanted organoids into mice for *in vivo* studies. M.H. prepared breast tumors from patients for primary cell experiments. A.K. and P.W. performed image analysis. T.M., M.H., I.M.S., and T.L.W., edited the manuscript. M.H.L. and D.W. wrote the manuscript with input from G.C.R., P.W., and I.M.S.

## Corresponding Author

Correspondence to Denis Wirtz

