## Supplemental Figures and captions for "Multi-compartment tumor organoids"

### Supplementary Figures

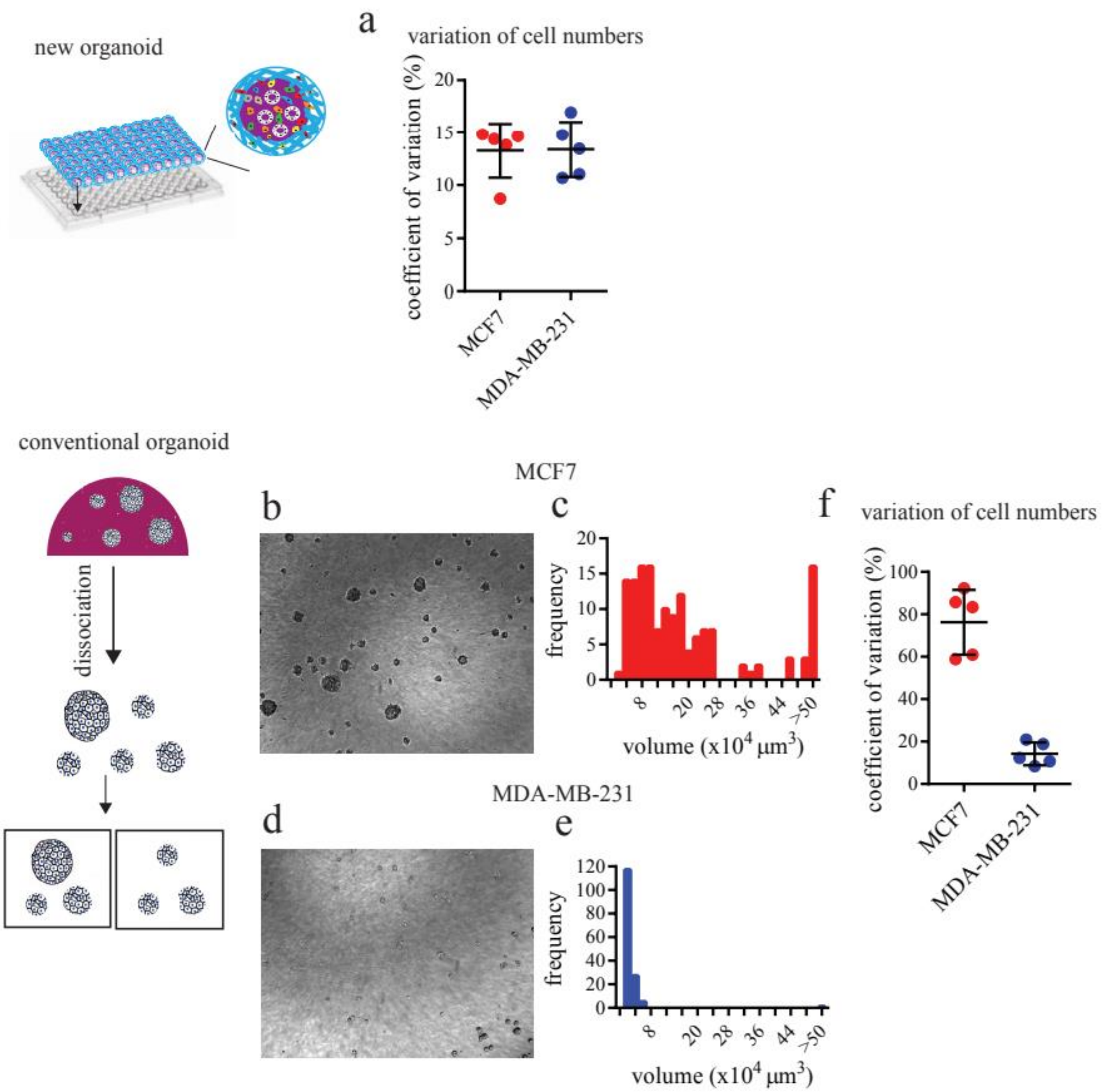


**Supplementary Figure 1**. **New organoids allow for high consistency in cell numbers per organoid.**

**a,** Coefficient of variation ± SEM in the mean cell number per organoid for the new two-compartment organoids containing either MCF7 or MDA-MB-231 breast cancer cells in their core. A total of 140 two-compartment organoids was assessed for 5 biological repeats for each cell line. **b**, Representative phase-contrast image and **c,** volume distribution of conventional organoids containing MCF7 cells in Matrigel. **d**, Representative phase-contrast image and **e,** volume distribution of conventional organoids containing MDA-MB-231 cells in Matrigel. **f,** Coefficient of variation ± SEM in the mean cell number per organoid for conventional organoids containing either MCF7 or MDA-MB-231 cells in Matrigel. A total of 140 conventional organoids was assessed for 5 biological repeats for each cell line.

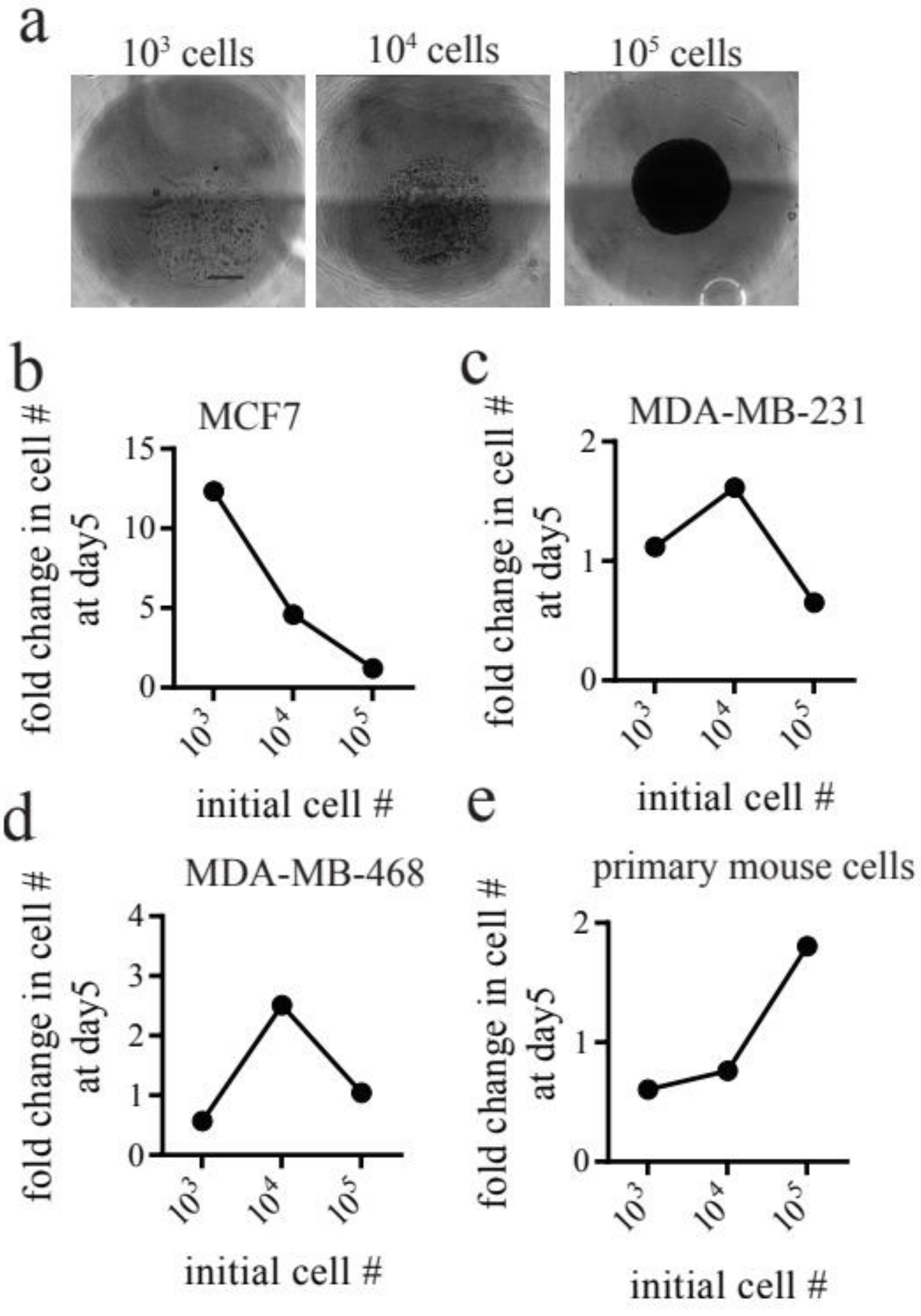

**Supplementary Figure 2**. **Growth of cancer cells in the new organoids can be optimized by adjusting seeding cell density.**

The number of cancer cells seeded in the core of two-compartment modulates the growth pattern of the organoid. **a,** Representative phase-contrast images of new organoids with seeding cell numbers ranging from $10^4$ to $10^6$ cells imaged at day 2. Change in the number of cells per organoid measured at day 5 after seeding for new organoids containing either **b,** MCF7, **c,** MDA-MB-231, **d,** MDA-MB468 breast cancer cells or **e,** primary mouse cells isolated from the uterus of mouse models of endometrioid carcinoma.

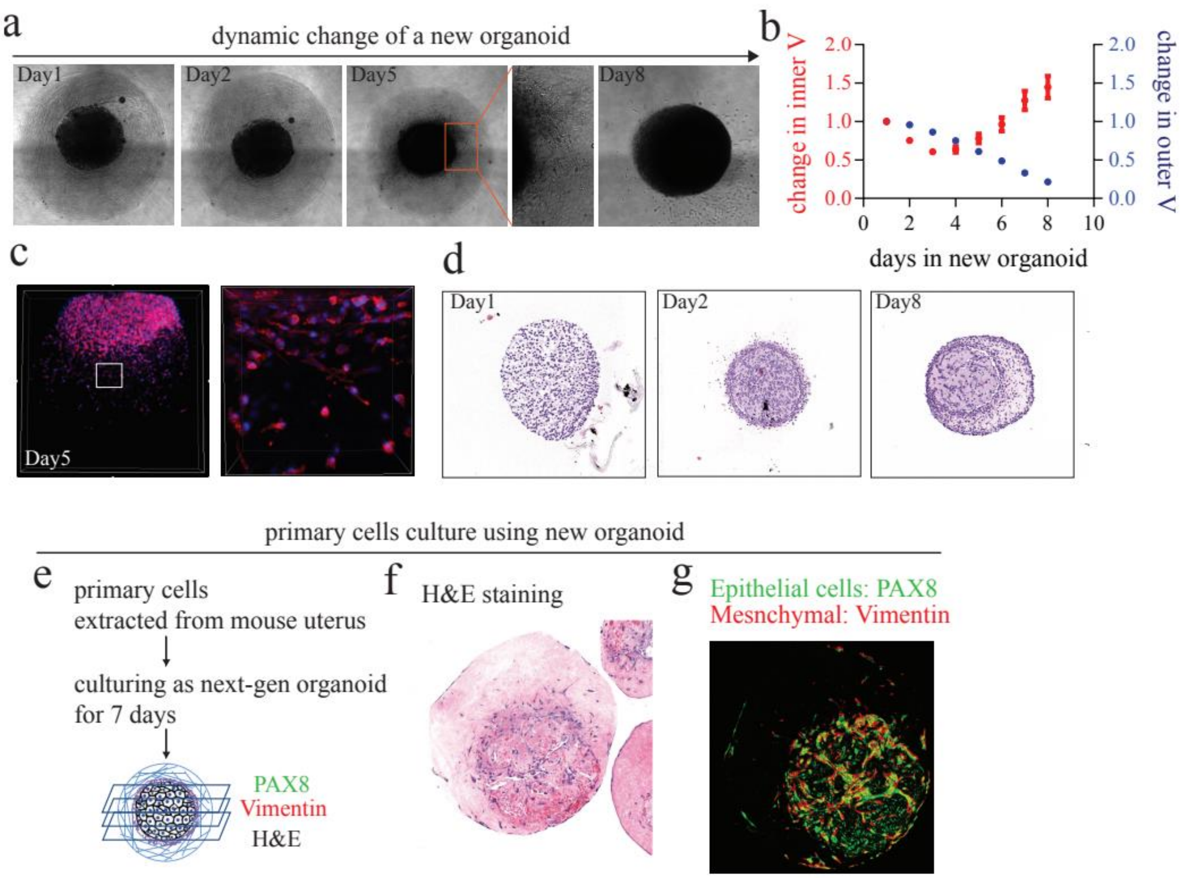


**Supplementary Figure 3. Evolution of new organoids imaged under different microscopies.** Similarly to conventional organoids, two-compartment organoids can be subjected to different types of light and scanning microscopy. **a,** Time-dependent morphological changes and **b,** associated time-dependent volumetric changes of the inner tumor Matrigel core and the surrounding outer collagen I compartment of two-compartment MDA-MB-231 organoids via bright-field microscopy. **c,** Immunofluorescence confocal images of a two-compartment organoid in which the MDA-MB-231 cells were labeled with F-actin (red) and nuclear DNA (blue). **d,** Hematoxylin and eosin (H&E) stained cross-sections of two-compartment MDA-MB-231 organoids as a function of days in culture. **e,** Schematic of a two-compartment organoid containing primary mouse cells subjected to sectioning, immunohistochemistry (IHC) or H&E staining, and scanning. **f and g,** Representative H&E image (f) and IHC image for a slide stained for the PAX8 protein and cytoskeletal protein vimentin (g) for two-compartment organoids containing cells dissociated from the uterus of mouse models of endometrioid carcinoma after 7 days in culture. Scale bar, 200 μm.

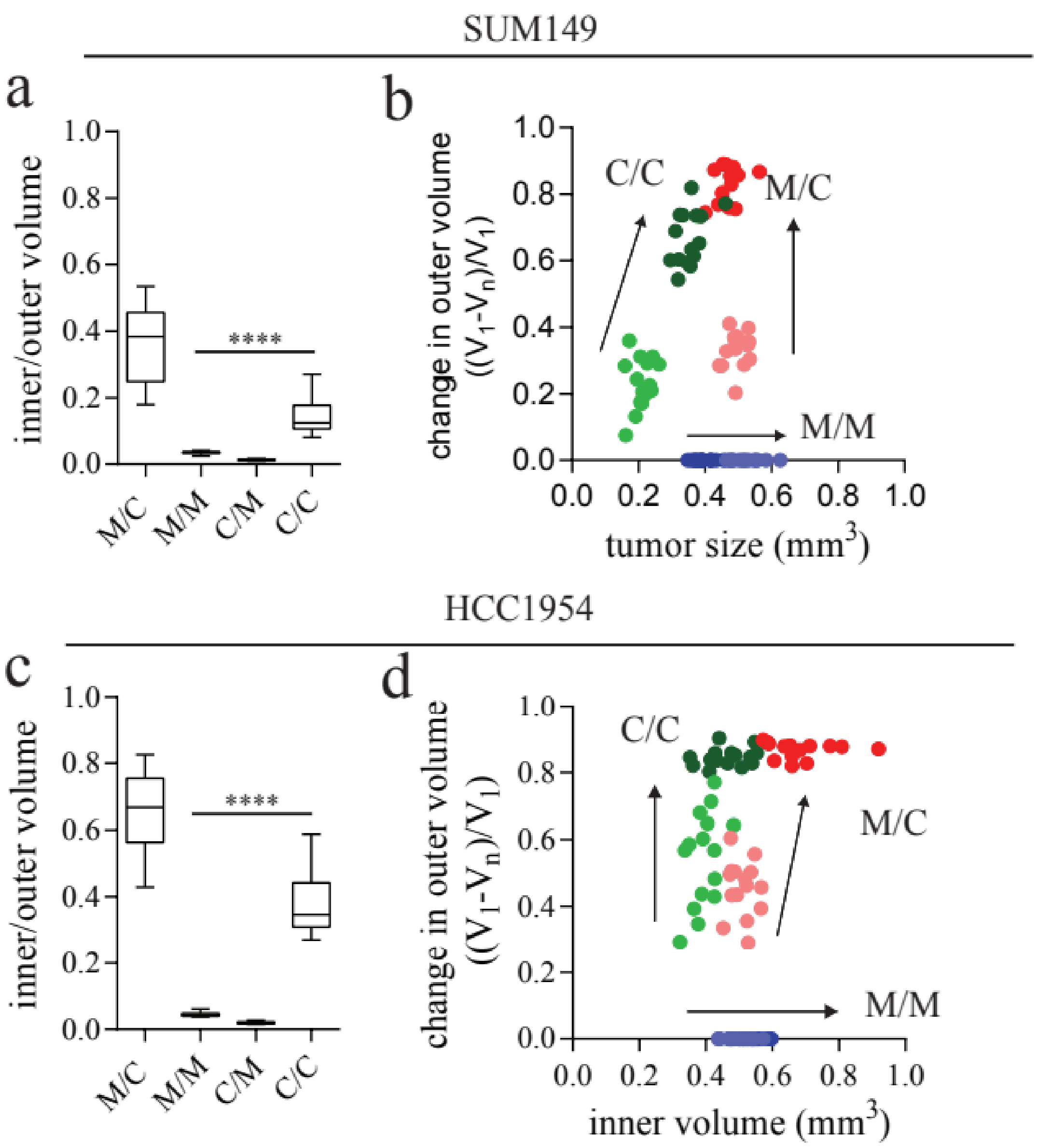

**Supplementary Figure 4**. **Dynamic changes in volume of two-compartment organoids containing different types of cancer cells.**

Bar graphs representing the mean ± SEM of the dynamic changes in two-compartment organoid morphology indicated by the ratio of the volume of the inner core and the outer compartment for new organoids containing **d**, SUM149 and **c,** HCC1954 breast cancer cells. Shrinkage of the outer compartment and expansion of the tumor core between day3 and day5 for new organoids containing **b**, SUM149 and **d,** HCC1954 breast cancer cells. Fifteen organoids encompassing three biological repeats were tested for each measurement.

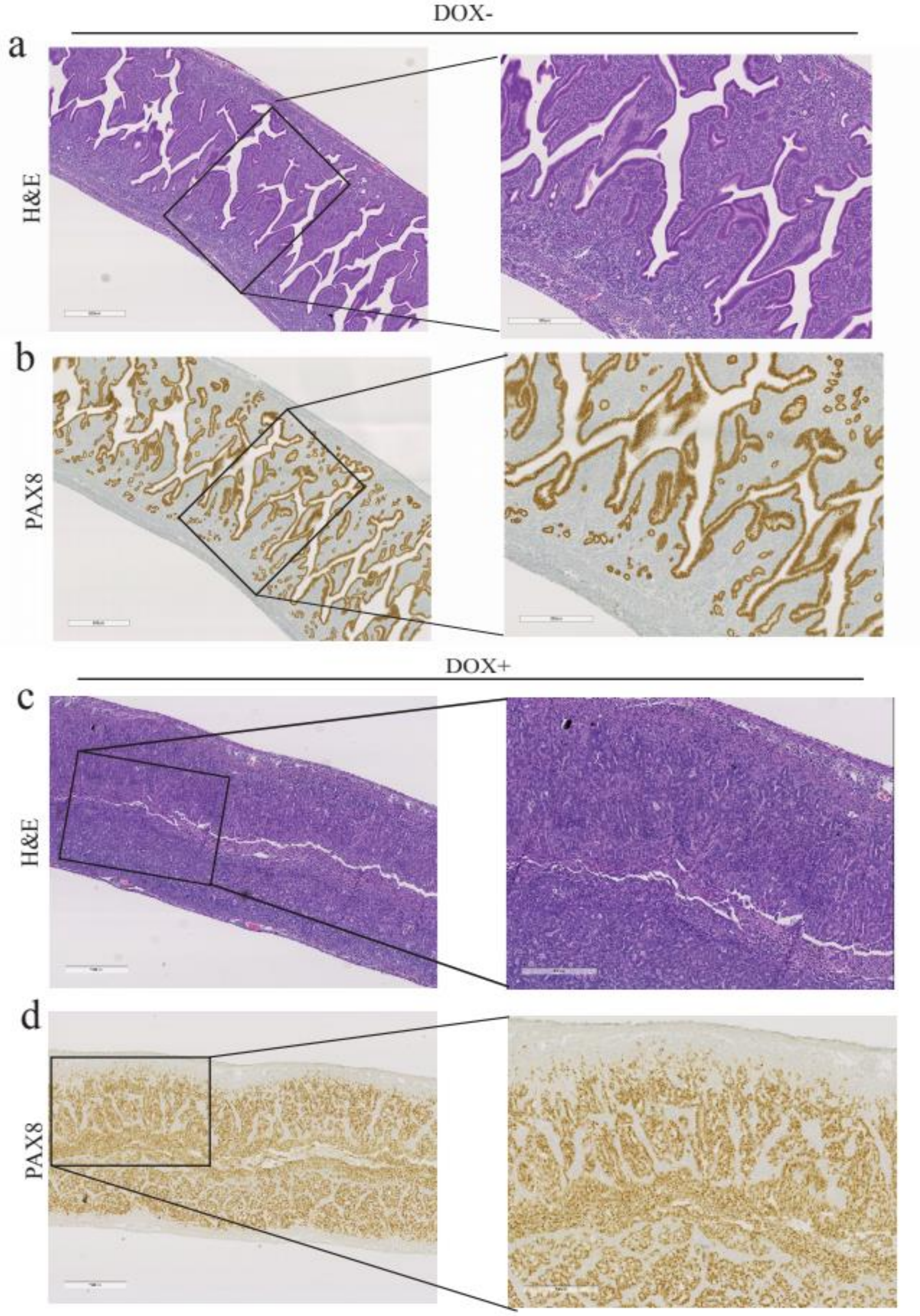
DOX-
a
H&E
b
PAX8
DOX+
c
H&E
d
PAX8

**Supplementary Figure 5**. **Organoids from doxycycline-inducible *ARID1A* and *PTEN* knockout endometrial cancer mouse models.** Representative (**a** and **c**) H&E images and (**b** and **d**) immunohistochemistry images stained with PAX8 of the uteruses of mouse models of endometrioid carcinoma treated with either (**a** and **b**) vehicle or (**c** and **d**) doxycycline.

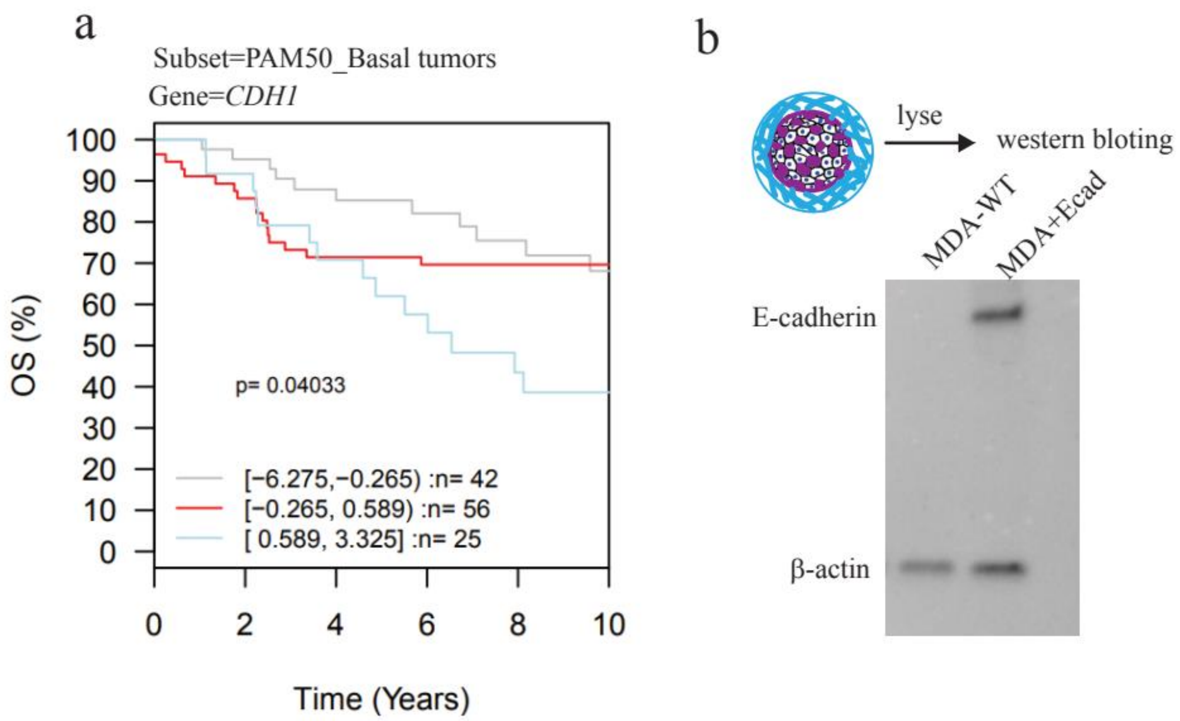


**Supplementary Figure 6**. **Effect of E-cadherin expression on tumor organoid progression.**

**a,** Kaplan-Meier curve showing the correlation between E-cadherin (*CDH1*) expression and overall survival (OS) in primary basal breast tumors characterized by PAM50 genes in the GOBO database[55]. **b,** Representative Western blot showing the stable expression of E-cadherin in MDA-MB-231 cells used in the organoids transduced using lentiviral system.

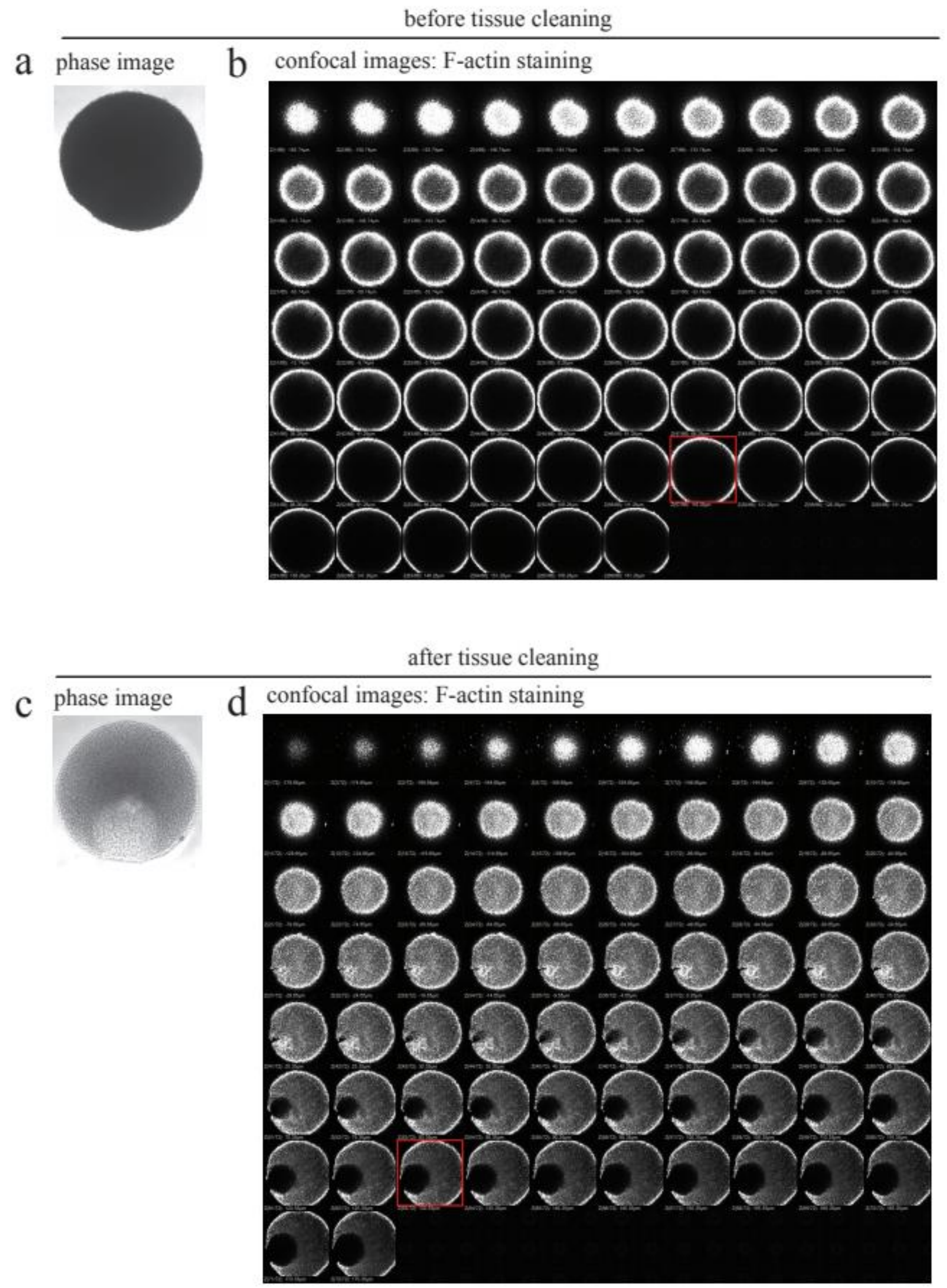


**Supplementary Figure 7**. **Tissue clearing of two-compartment organoids.**

MDA-MB-231 cells grew in two-compartment organoids for 7 days. The cells were then fixed and labeled with F-actin. **a,** Representative phase-contrast image and **b,** confocal scanning images of the new organoid before application of the tissue-clearing reagent. **c,** Representative phase-contrast image and **d,** confocal scanning images of the new organoid after application of the tissue-clearing reagent.

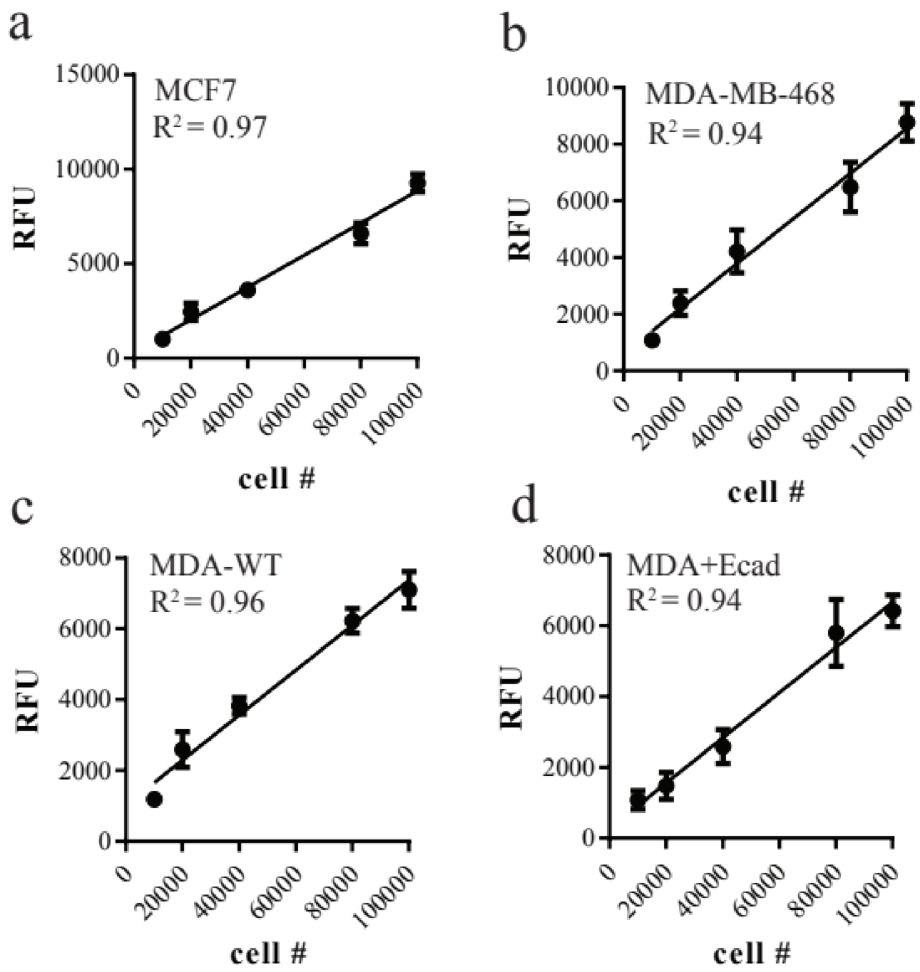


**Supplementary Figure 8**. **Measurement of cell number in two-compartment organoids.**

The measurement of the number of cells in an organoid is measured using standardization curves. These curves show a linear correlation between the controlled cell numbers seeded in the core of an organoid and assessed within 6 h and the relative fluorescence units (RFU) of Prestoblue for two-compartment organoids containing. Standardization curves for **a,** MCF7 breast cancer cells, **b,** MDA-MB-468 breast cancer cells, **c,** (scramble control) MDA-MB-231 breast cancer cells (MDA-WT), and **d,** MDA-MB-231 cells stably expressing E-cadherin (MDA+Ecad).
